# Mitochondrial Complex I activity is inhibited by changes in the abundance of phosphatidylinositol and phosphatidylserine in wheat plants exposed to high temperatures

**DOI:** 10.1101/2023.02.05.527222

**Authors:** Aygul Malone, Thusitha W. Rupasinghe, Ute Roessner, Nicolas L. Taylor

## Abstract

Identifying the molecular basis of thermotolerance in crops is becoming increasingly important with the changing climatic conditions that challenge future food security. Sustaining cellular energy production under heat stress is vital in maintaining an uninterrupted growth cycle, and thus the mitochondria is instrumental in facilitating the overall heat-tolerance of a crop plant. Using targeted mass spectrometry, the changes in abundance of the lipo-protein network in mitochondrial membranes following a short episode of extremely high temperature were analysed in two wheat cultivars of differing thermotolerance. The results indicated that membrane lipids remodel in favour of shorter fatty acyl tails, and an increase in the abundance of phosphatidylinositol, while specific to the heat-tolerant cultivar was an increase in the abundance of phosphatidylserine. The differences between the lipid profiles of the two cultivars is a likely explanation for the decrease in Complex I NADH dehydrogenase activity in the heat-sensitive cultivar. Further metabolite analysis by LC-MS revealed malate accumulation, indicating that the disruption in Complex I activity impacts the catabolism of reducing equivalents. The measured increase in the total amount of phosphatidylserine in the heat-tolerant cultivar suggests a potential role in conveying thermotolerance for this minor membrane constituent, and highlights that a focus on membrane lipid composition during thermal stress will be essential for the breeding of future heat tolerant crops.

**Summary:**

- We evaluated changes to the lipo-protein network of wheat mitochondria of differing heat tolerance in response to heat shock.
- Using targeted mass spectrometry, candidate transitions were selected to quantify changes in membrane lipids and the embedded protein components of the electron transport chain, which play a vital role in maintaining respiration.
- A significant increase in phosphatidylserine was exclusive to the mitochondria of the heat-tolerant wheat cultivar. In the absence of this, the heat-sensitive cultivar displayed a reduced Complex I activity.
- The minor membrane constituent phosphatidylserine plays a role in conveying thermotolerance, making this membrane lipid a focal point for the breeding of future heat tolerant crops.

## Introduction

As the demand for wheat and wheat products increases, the current grain yield needs to increase by 60% by 2050 to sustain the needs of the growing global population (Peña-Bautista et al. 2017). Higher atmospheric temperature accelerates the rate of growth, bringing about early leaf senescence and truncating the time available for grain formation, as well as the abortion of grain formation (Asseng et al. 2011). Therefore, the projected climate change scenario, with its accompanying increase in the frequency, intensity and duration of heat shock events, poses a significant challenge to achieving increased wheat yields to ensure global food security (www.bom.gov.au (2014); (Barlow et al. 2015)).

Grain formation is an energy expensive process, relying on the biomass assimilated during the vegetative phase as the energy source, and as such, higher biomass often correlates with higher grain yield (Ferguson et al. 2021, Sharma 1993, Wilson et al. 1982, Hauben et al. 2009). However, this means that a heat shock event occurring at the vegetative growth stage has a downstream impact on yield (Janni et al. 2020). To further compound this issue, the duration of grain filling is 4% shorter in heat-sensitive wheat cultivars in response to warmer temperatures, compared to the heat-tolerant counterparts (Stone et al. 1995, Morison et al. 1999). This highlights the need to effectively identify those wheat cultivars that can adapt to, and withstand, the changing climate.

The trait of heat tolerance is defined as the ability to maintain an uninterrupted growth cycle in the presence of heat stress (Janni et al. 2020). Given that the energy required for plant growth and repair is synthesised in the inner mitochondrial membrane (IMM) via the electron transport chain (ETC), it has been proposed that thermotolerance of the mitochondria leads to the acquiring of thermotolerance of the whole plant (Sanmiya et al. 2004). With the close interaction between the membrane proteins and the lipid bilayer, it is no surprise that the structure and activity of the components of the ETC are highly dependent on the presence of specific phospholipids in the membrane (Bogdanov et al. 1996, Schägger et al. 2000, David et al. 2002, Sazanov et al. 2003, Sharpley et al. 2006, Bogdanov et al. 2008, Pineau et al. 2013, Petereit et al. 2017, van Meer et al. 2008).

Membrane lipids undergo phase transition at specific transition temperature T_m_, changing the physical state of the bilayer (Marsh 2010, Raudino et al. 2011). In order maintain the metabolic reactions that are dependent on membrane architecture, plants modify their membrane lipid composition in response to temperature to preserve membrane integrity and lipo-protein interactions (Chen et al. 1982, Neidleman 1987, Hazel 1995, Vanlerberghe 2013). Previous work has suggested that tolerant wheat cultivars have a higher capacity for rapid lipid remodeling during temperature stress when compared to their sensitive counterparts (Chen et al. 1982, Narayanan et al. 2016). Here, we sought to compare the impact of short-term heat shock on the changes in the lipo-protein network of two wheat cultivars, heat tolerant cv. Mace (Mace) and heat sensitive cv. EGA2248 (EGA2248) (Collins 2015; Lyon 2017; Trethowan 2020). Taken together, we propose that the key differences in the lipid remodeling during heat shock convey thermotolerance through Complex I activity, perpetuating energy production through the ETC towards biomass accumulation, which can subsequently be drawn upon during grain formation.

## Materials and Methods

### Biomass measurements

Mace (Australian Grain Technologies) and EGA2248 (InterGrain) seeds were pre-soaked overnight in water and allowed to germinate at room temperature for five days. Viable seedlings were transferred into individual pots filled with 3:1:1 peat: vermiculite: pearlite, and grown in plant growth chamber under controlled conditions of 24 °C day-time, 20 °C night-time temperature, 65 % humidity, 300 μmol light for 16 h, and a 8 h dark cycle (optimal growth conditions, OT). Heat shock (HT) was performed by increasing the temperature to 36 °C day-time, 31 °C night-time for two days on day 12. 14-day old seedlings were harvested, and the total height recorded. Lyophilized seedlings were weighed for dry mass measurements. The mean was calculated for each variety under each growth condition, and outliers greater than one standard deviation were excluded from further analysis. The number of biological replicates was n=≥10 for per variety, per growth condition.

### Isolation of mitochondria from wheat seedlings

Rehydrated seeds were scattered over a 3:1:1 peat:vermiculite:pearlite mixture, and grown in a controlled environment chamber under controlled OT or HT conditions as described previously. The trays were watered daily to avoid water limitation. Seventy to eighty grams of fresh wheat shoots were harvested following 14 days of growth, and mitochondria were isolated using PVP-Percoll gradient centrifugation following the method described by Kerbler & Taylor (2017). The mitochondrial pellet was stored at −80 °C. The protein concentration of the mitochondrial extract was determined using a Bradford Assay with BSA as a standard (Bradford 1976).

### Phospholipid profiling by LC-QqQ mass spectrometry

Lipids were extracted from isolated mitochondria using methyl-*tert-*butyl ether (MTBE), as described by Matyash et al. (2008). Lipids were extracted from 100 μg of mitochondrial protein, with four biological replicates for each treatment condition. Dried lipid aliquots were stored at −80°C.

Dried lipid aliquots were resuspended in 100 μL of loading buffer (50:50 (v/v) butanol:methanol containing 10 mM ammonium formate). Ten μL was injected onto an Poroshell 150 × 2.1mm C18-EC column attached to an 1100 series LC coupled to an Agilent 6470 QqQ mass spectrometer (Agilent Technologies, Santa Clara). Column temperature was set at 60 °C. The chromatography used a mobile phase A of 20:30:50 (v/v) isopropanol: acetonitrile: MS-grade water, containing 10 mM ammonium formate, and lipids were eluted by a by increasing mobile phase B (90:9:1 (v/v) isopropanol: acetonitrile: MS-grade water, containing 10mM ammonium formate) from 10-50 % over the first 4 minutes, 50-65 % gradient for 11 minutes, increasing to 100 % B for three minutes, and a final hold at 100 % for one minute. Flow rate was set to 200 μL/min. Drying gas flow was set to 13 L/min, nebulizer pressure at 40 psi and source temperature at 250 °C. A pooled biological quality control (PBQC) sample containing 10 μL of each biological replicate was used to compile the phospholipid target list for selected reaction monitoring (SRM), and injected at every 10^th^ sample to monitor the LC-MS sequence runs. The phospholipid target list is detailed in Supplementary Table 1. Lipids were quantified as described by Natera et.al (2016), and expressed in nmol mg^−1^ protein, ± SEM, n=4.

### Detection and relative quantification of cardiolipin

Probable cardiolipin species were first detected as singly charged MS1 ions in negative ion mode, following the chromatography conditions described previously. Verified cardiolipin species were included in the QqQ method (Supplementary Table 2).

### Respiratory activity assays and complex abundance

Electron transfer chain complex activity assays were carried out in a 96-well flat-bottomed plate, in a final volume of 200 μL and changes in absorbance was measured at 30 °C (BMG Labtech Spectrostar Nano). The Beer-Lambert equation (A=ɛcl) was rearranged to determine the molar concentration, with the path length 5.88 mm. The reaction rate was expressed in molar concentration min^−1^mg^−1^ mitochondrial protein. The assay conditions are adapted from Ivanova *et al*. (2019).

The relative abundance of respiratory complex proteins was measured using SRM mass spectrometry, adapted from Taylor et al. (2014). Acetone precipitated proteins from 100 μg of mitochondrial protein were reduced, alkylated and digested as described by Taylor et al. (2014). Peptides were solid phase extracted using a C18 Macrospin column (The Nest Group) and a positive pressure manifold, following the manufacturers’ specifications, and the eluate dried under vacuum. The resulting pellet was resuspended in 50 μL of 5 % (v/v) acetonitrile with 0.01 % (v/v) formic acid, to a final concentration of 2 μg μL^−1^, and 5 μL injected into an Infinity HPLC 1300 coupled to an 6495 QqQ mass spectrometer (Agilent Technologies, Santa Clara). Peptides were loaded onto a C18 120 Å 2.7 μm 2.1 × 250 mm peptide mapping column held at 60 °C. The stationary phase was HPLC grade 100% (v/v) Water + 0.1 % (v/v) formic acid, the mobile phase was 100% acetonitrile with 0.1 % (v/v) formic acid. Peptides were eluted directly into the mass spectrometer during a 40 minute, 3-35 % gradient. The mass spectrometer was run in positive ion mode, with a gas temperature of 250 °C at a flow rate of 15 L min^−1^. The maximum dwell time was 165.20 ms. Collision energy was optimized by the predicate values by Skyline. A pooled biological quality control (PBQC) sample, containing 5 μL of each biological replicate was injected at every 10^th^ sample to monitor the LC-MS sequence runs.

SRM transitions were selected from an in-house database of wheat mitochondrial proteins developed from the TGAT 1.0 genome sequence. Eighty-four peptides passed the criteria for matching the spectral library with one qualifier and 2 quantifiers, were detected in the pooled biological quality control (PBQC) sample and were included into a SRM list (Supplementary Table 3). Accession ID were updated by searching the peptides against IWGSC RefSeq 1.0 using https://wheatproteome.org/. All hits are listed in Supplementary Table 4. For convenience, the lowest genome ortholog is deemed a protein match, all other orthologs are also listed. Data was checked on Skyline (version 20.2.0.286). The area under the curve (AUC) from the 1^st^ ranking ion was used to determine relative of peptide abundance.

### BN-PAGE activity assay

Four hundred micrograms of pooled mitochondrial proteins were used for each variety and treatment. Mitochondrial protein pellet solubilisation, with BN-PAGE using a 4.5 %-16 % native gel, was carried out as described in Ivanova *et al*. (2019). The Complex I in-gel NADH-dehydrogenase activity stain was carried out as described by Sabar et al. (2005). The bands were identified by comparison to the results of Gazizova et al. (2020), who performed their assay on protein complexes isolated from wheat root mitochondria. The gel was rinsed thoroughly with double deionised water (DDI) prior to protein staining with colloidal Coomassie (Eubel et al. 2005). BN-PAGE band intensities were quantified using GelAnalyzer software (version 19.1, www.gelanalyzer.com).

### Quantification of TCA cycle intermediates by LC-QqQ-MS

Metabolites were extracted from approximately 30 mg of pre-ground, snap-frozen green tissue with 500 μL extraction medium (90% (v/v) methanol containing internal standard 8 mg/L adipic acid). The slurry was incubated at on a thermomixer set at 75 °C, 1200 rpm, for 20 minutes. Debris were pelleted by centrifuging at 20 000 x g for 3 minutes. One hundred microlitres of the supernatant was transferred to a clean tube, and dried by vacuum centrifugation. Samples were stored at −80 °C until derivatization with 3-nitrophenylhydrazine (3-NPH) as described by Li *et al*. (2021).

Standard curves for each of the metabolites, as well as the internal standard adipic acid, were constructed on Agilent Technologies MassHunter quantitative analysis for LC-MS (version B.07.01), with quadratic curve fit and origin ignore. Twenty μL was injected onto Phenomenex Kinetex XB-C18 column (50 × 2.1 mm, 5 μm) for the SRM detection of lower abundant metabolites pyruvate and α-ketoglutarate, with 1 μL injected for all other metabolites. Sample analysis by LC-QqQ-MS and the SRM transitions are described in detailed in Xuyen et al. 2021.

### Data analysis

Lipidomics data was inspected for quality and adjustments made to integrated peaks using Agilent MassHunter Quantitative Analysis software (version B07.01, 2008). For relative abundance assessment, the peak the peak area of samples was normalised by median normalization. One-way ANOVA and Tukey Honest Significant Difference (HSD) tests at a 5 % significance level were performed in *R* (R Core Team 2019). Figures were produced with the ggplot package (Wickham 2016). Outliers greater than 1 standard deviation were excluded from statistical analysis for complex activity assays. Peptide relative abundance was assessed by calculating z-scores, and plotting as a heat map (Kolde 2019).

## Results

### Heat sensitivity is evident in the vegetative growth stages of wheat

Australian wheat variety EGA2248 ranked last in an analysis of heat tolerance of 137 wheat genotypes (Collins 2015), while the more recently developed cultivar Mace has been observed to be highly heat tolerant, achieving high grain yield (Lyon 2017, Trethowan 2020). To determine whether the heat sensitivity reported at the grain filling stage is prevalent at the vegetative growth stage, the development of 14-day old seedlings was evaluated by measuring the dry mass and the height of the longest blade. The two wheat varieties performed differently to each other under normal growth conditions (OT), and with this in mind, the biomass measurements following heat shock (HT) were converted to a relative percentage of those recorded during normal growth conditions (OT), to allow for the evaluation of the proportion reduction of vegetative growth of the two wheat varieties. Following heat shock, the length, and dry mass of Mace was reduced to 90.05%, and 75.51% respectively, while for EGA2248, the length and dry mass were reduced to 81.86% and 58.73%, respectively (Figure 1 a, b). Heat shock has a significantly greater impact on the growth of EGA2248 than Mace, apparent by the larger percentage reduction in biomass measurements in this variety. Previously, EGA2248 and Mace cultivars were shown to differ in their heat tolerance, as determined by grain size (Collins 2015, Trethowan 2020). Here, the vegetative growth of EGA2248 is decreased to a significantly larger extent than that of Mace. This indicates that the sensitivity to heat shock is evident at vegetative growth.

**Figure 1.**
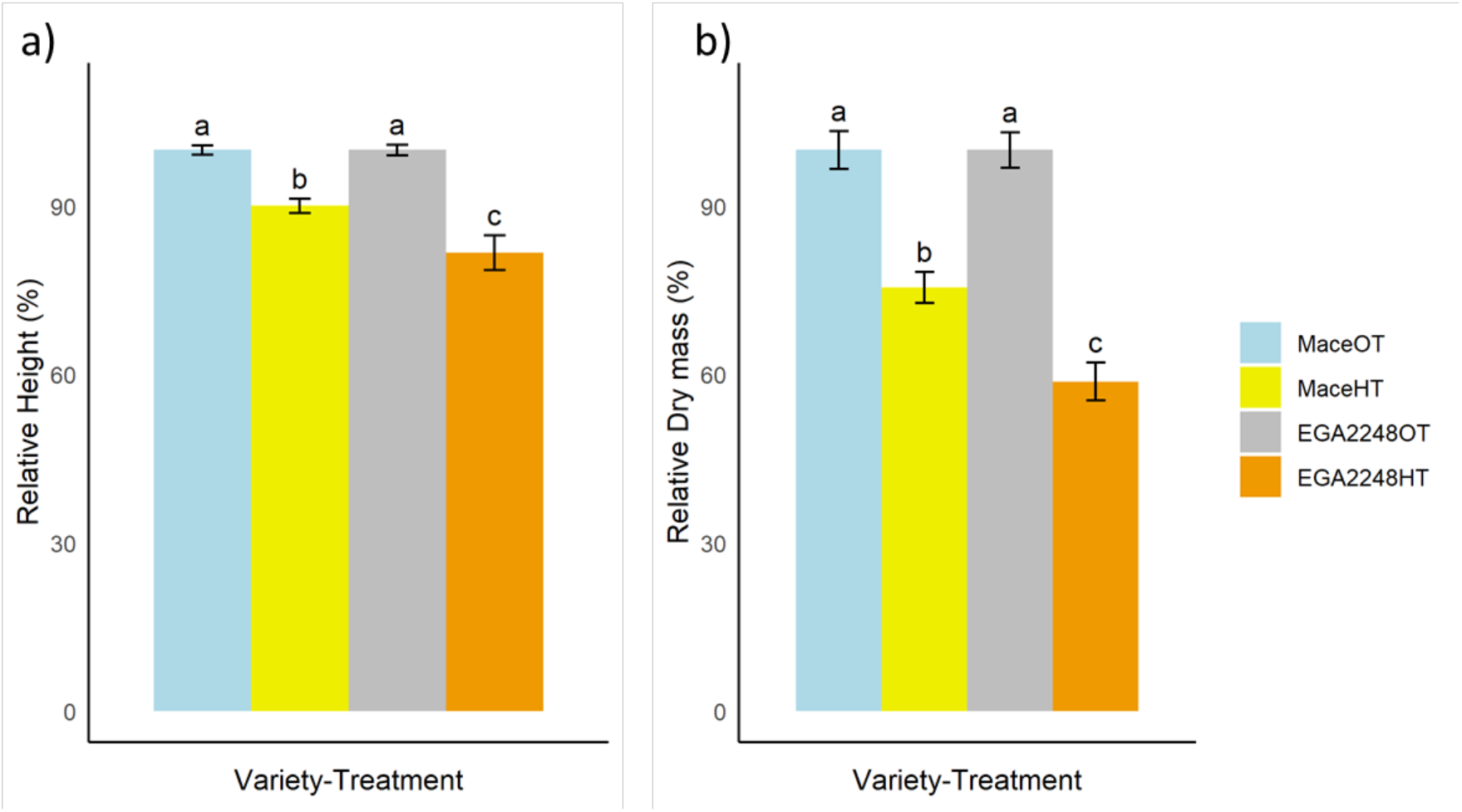
**a)** The decrease in the height and **b)** the dry mass of Mace and EGA2248 seedlings following heat treatment (HT), expressed as a relative percentage of the measurements obtained following growth at optimal temperature (OT). The impact of high temperature was greater in the heat-sensitive wheat variety, EGA2248, noted by the larger reduction in biomass accumulation (**a-c;** two-way ANOVA with repeated measures p ≤ 0.05). Values are the mean of n≥10 biological replicates (± SEM).

### Mitochondrial lipid remodelling following heat shock

It has been noted that heat tolerant wheat cultivars display a greater capacity for lipid remodelling (Narayanan et al. 2016), and here we sought to determine if lipid remodelling of the mitochondria confers heat-tolerance, and thus the perpetuation of the growth cycle, which was demonstrated by biomass accumulation. One-hundred and forty-eight lipid species across ten lipid classes were detected, with the absolute quantities, expressed as the nmol of lipid per mg of mitochondrial protein (nmol of lipid mg^−1^ protein) summarised in Figure 2a (Supplementary Table 5).

**Figure 2:**
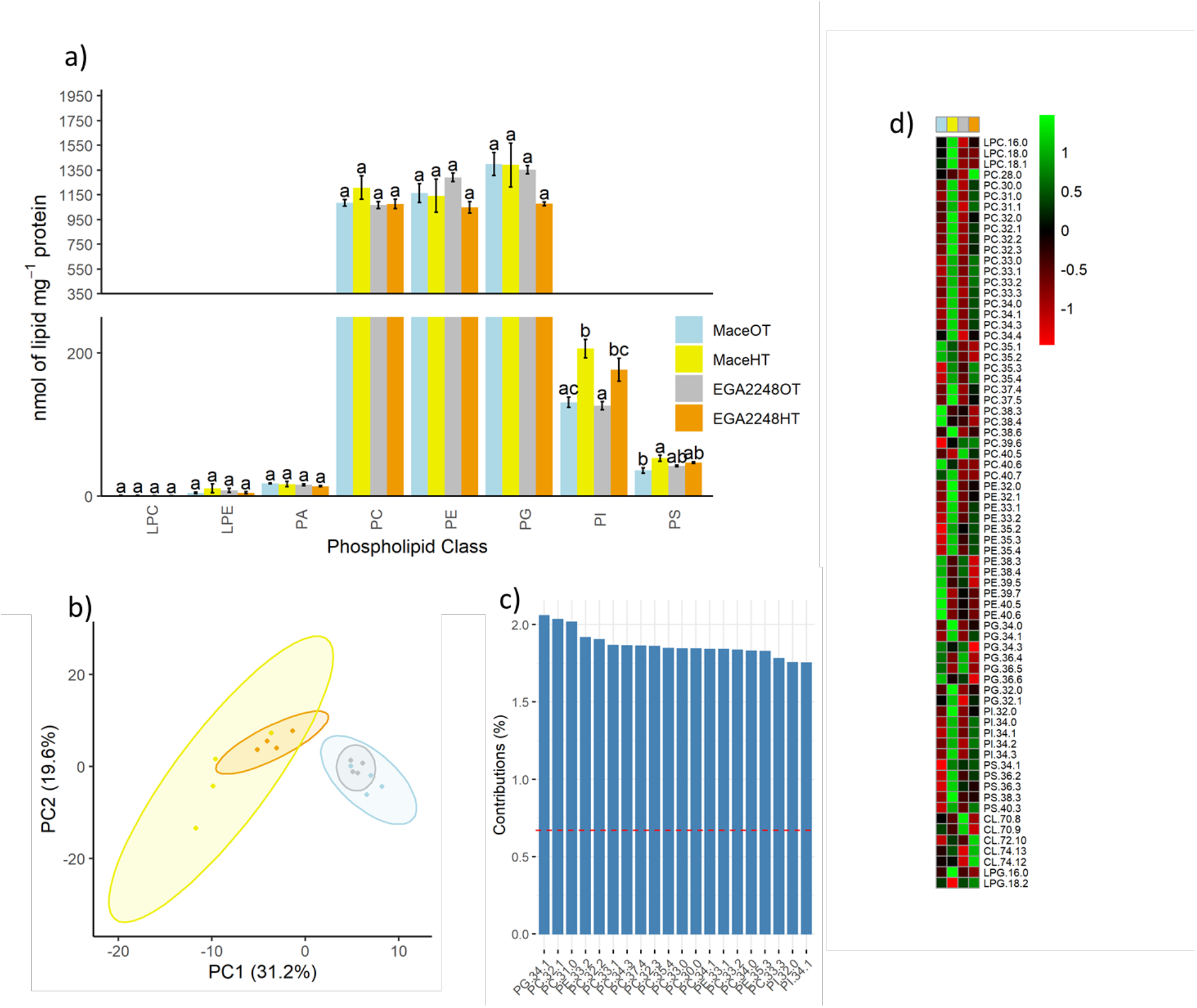
**a)** Absolute quantitation (nmol of lipid mg^−1^ mitochondrial protein) of lipid species quantified in isolated mitochondria obtained from OT and HT treated wheat seedlings. Values are derived from 4 biological replicates (**a-c;** two-way ANOVA with repeated measures p=≤ 0.05), ± SEM. **b)** The 148 lipid species across 10 lipid classes cluster in their expression according to growth temperature, the graphed points represent individual biological replicates. **c)** Top 20 lipid species contributing to principal component 1 include are phospholipids with shorter and odd-chain fatty acyl tails **d)** Heatmap of the 70 significantly changing lipid species. Values are derived from 4 biological replicates (**a-c;** two-way ANOVA with repeated measures p=≤ 0.05), ± SEM.

The major phospholipids of wheat mitochondrial membranes are PC and PE, accounting for ~40-45% and ~25-30% of the membrane respectively (Mårtensson et al. 2017), while mitochondrial membranes also contain CL (~15%) and PG (~5%) (Schwertner et al. 1973, Flis et al. 2013, Schenkel et al. 2014, Mårtensson et al. 2017, Liu et al. 2022). The total quantitation (nmol of lipid mg^−1^ mitochondrial protein) of lipid species in isolated mitochondria obtained from wheat seedlings demonstrate that the main membrane lipid constituents in both varieties, as expected, are PC and PE, accounting for 27-33% of membrane lipids, and that this does not differ significantly between the two varieties or with temperature regime. The total amount of PG, at 31-37%, is greater than previously detected in cauliflower and Arabidopsis mitochondria (Schwertner et al. 1973, Liu et al. 2022). PA is an intermediate in the biosynthesis of phospholipids and by itself is a minor constituent membranes (Thakur et al. 2019), and the abundance of PA detected did not significantly differ between the two wheat varieties. In this study, lyso-phospholipids LPC, LPE and LPG were detected, with the abundance of LPE 10 times greater than the other LPCs.

When comparing the mitochondrial lipid profiles of the two wheat cultivars grown at OT and HT, two distinct clusters from the principal component analysis are evident (Figure 2b). Thirty one percent (31.2%) of the differences in the two clusters can be explained by lipids that have a large influence on principal component 1 (PC1). The highest contributing lipid species to PC1 are phospholipids containing an odd-chain fatty acyl side chain (Figure 2c). Based on this, the length of the acyl tail appears to have the largest influence on the separation of the clusters. The total amount of PI increased by ~50% during HT. Also notable is the increase in the total amount of PS in the heat tolerant variety, Mace, during HT. In Mace, PS increased from 35.56±3.54 to 52.62±4.10 nmol of lipid mg^−1^ protein, whereas the quantity of PS in EGA2248 did not change significantly with a change from 41.87 ± 1.11 to 46.84 ± 1.10 nmol of lipid mg^−1^ protein after heat treatment.

One-way ANOVA with Tukeys Honestly Significant Difference (HSD) test indicated significant differences in the expression of 70 lipid species following exposure to high temperatures. Z-scores for the lipid features significantly impacted in their expression were plotted as a heat map to allow for the visualization of the membrane lipid remodelling (Figure 2d). There is an indication that the lipid remodeling varies between the two wheat cultivars. The largest and smallest z-scores on the heat map stem from the abundance of lipids in Mace, and this indicates a more effective and rapid lipid remodeling in response to HT. Notable remodeling exclusive to the heat tolerant cultivar following HT are the increase in the abundance of LPC 18:0 and 18:1, and the trend towards PE with longer acyl chain length of 38, 39 and 40 carbons to decrease in abundance under HT, with PE 40:5 and 40:6 significantly decreasing in Mace during HT. The increase in PS in Mace can be attributed to the accumulation of longer-chain PS species PS 38:5, 42:3. The quantity of PG 32:0, 32:1, 34:0 and 34:1 increased during HT while PG 36:4 and 36:5 decreased. This pattern of remodeling follows that observed in PC, PE and PI, favouring shorter chain acyl tails. The pattern of CL expression was markedly different between the two wheat varieties. The relative quantities of CL 70:8, 70:9 decreased in EGA2248 during HT exposure. The most abundant of the CLs detected, 72:10, increases in EGA2248 during HT, as do 74:12 and 74:13. In the current study, numerous lipids containing odd-chain fatty acyl side chains were detected. The abundance of PC and PE with odd-chain fatty acyl side chains significantly increased during HT. Interestingly, no odd-chain fatty acyl side chains were detected from the other lipid classes. PC 31:0, 31:1, 33:1, 33:2, 33:3, 35:3, 35:4, 37:4, and PE 33:1 all increased two-fold during HT. PE 39:5 and 39:7 expression decreased two-fold during HT. There were some differences in the expression of odd-chain phospholipids between the two varieties, with PE 35:2, 35:3, 35:4 increasing two-fold during HT in Mace only.

### Decrease in NADH oxidation in the heat-sensitive wheat variety EGA2248 following heat stress is unrelated to protein abundance

With growing evidence of the role of membrane lipids in maintaining protein structure and regulating activity (Bogdanov et al. 2008), we aimed to discern the effect of the lipid remodelling on the mitochondrial lipo-protein network (Figure 3 a, b). To do so, the specific activity of respiratory complexes I-IV was measured.

**Figure 3.**
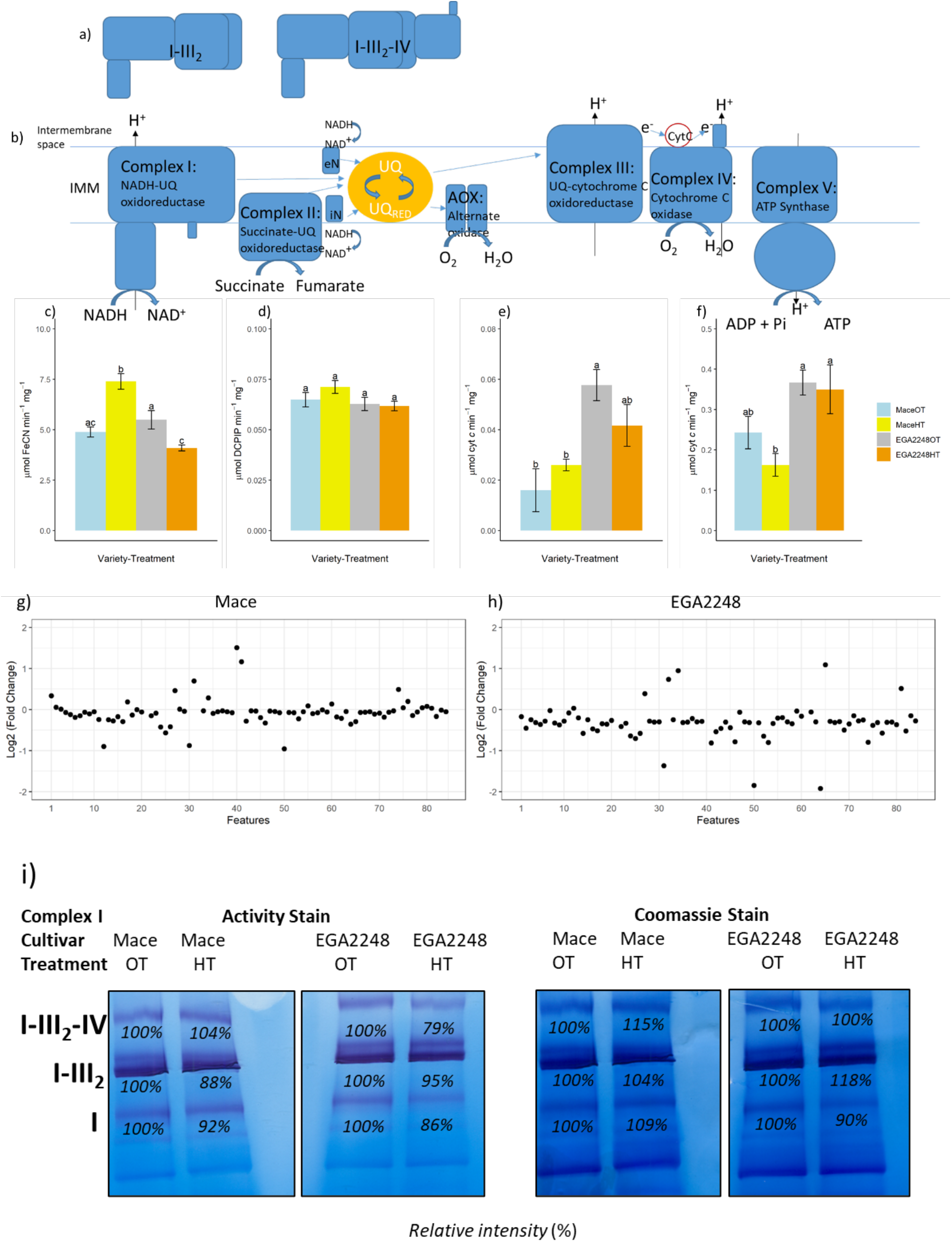
**a)** Schematic diagram of the assembly of respiratory complexes into supercomplexes and **b)** the arrangement of complexes to form the electron transport chain, localized in the inner mitochondrial membrane **c)** Specific activity of Complex I, determined as μmol K_3_Fe(CN)_6_ reduced min^− 1^ per mg mitochondrial protein (μmol FeCN min^−1^mg^−1^). Values are the mean of n≥5 biological replicates (± SEM). **d)** Specific activity of Complex II, determined as μmol DCPIP reduced min^−1^ mg^−1^ mitochondrial protein (μmol DCPIP min^−1^ mg^−1^). Values are the mean of ≥4 biological replicates (± SEM). **e)** Specific activity of Complex III, determined as μmol cyt *c* reduced min^−1^ mg^−1^ mitochondrial protein (μmol cyt *c* min^−1^mg^−1^). Values are the mean of ≥3 biological replicates (± SEM). **f)** Specific activity of Complex IV, determined as μmol cyt *c* oxidised min^−1^ per mg mitochondrial protein (μmol cyt *c* min^−1^mg^−1^). Values are the mean of ≥5 biological replicates (± SEM). **(c-f)** two-way ANOVA with repeated measures p=≤ 0.05) **g)** Changes in the abundance of ETC proteins between OT and HT in Mace and **h)** EGA2248. Proteins were quantified using selective reaction monitoring mass spectrometry. **i)** ETC complexes isolated by BN-PAGE from the mitochondria obtained from OT and HT treated wheat seedlings. Complex I and supercomplexes I-III2-IV and I-III2 are indicated by arrows. Complex I activity staining by nitro blue tetrazolium (NBT), showing a decrease of EGA12248 Complex I activity and in the supercomplex I-III2-IV following heat shock, and Coomassie staining showing that the abundance of Complex I following heat shock does not increase in EGA2248. This suggests that the complex is not dissociating from the supercomplex assembly.

For wheat grown at OT, the specific activity of NADH dehydrogenases did not differ significantly between Mace and EGA2248, determined as 4.89 ± 0.25 and 5.49 ± 0.46 μmol min^−1^mg^−1^ protein, respectively (Figure 3c). These values are in-line with previously reported specific activity of NADH dehydrogenases from plants (Soole et al. 1992, Serge et al. 1993, Lin et al. 1995). When subjected to HT, the activity from mitochondria isolated from Mace increased significantly to 7.39 ± 0.39 μmol min^− 1^ mg^−1^ protein, whereas in the heat sensitive wheat EGA2248 there was a significant decrease in specific activity, to 4.09 ± 0.15 μmol min^−1^ mg^−1^ protein. As NADH was the substrate used in the assay, this activity is the NADH oxidation from both Complex I and type II NADH dehydrogenases. Type II NADH dehydrogenases are a simplified form of Complex I as they have an NADH binding site and one flavoprotein redox centre, but only consist of two subunits (Soole et al. 1995). They are anchored to the membrane through the C-terminus side of a homodimer (Feng et al. 2012). In yeast, the dimerization forms a membrane anchoring domain, which is required for protein function, with structural studies indicating a hydrophobic surface, as well as positively charged patches which interact with the phosphate groups of membrane phospholipids that directs the localisation of the protein to the surface (Feng et al. 2012). A possible mechanism for the enhanced activity is that the changes in the lipid profile of Mace during HT supports the dimerization and activation of type II NADH dehydrogenases, which is reflected in the increase in specific activity of NADH oxidation of Mace during HT.

The activity of Complex II from isolated mitochondria grown at OT was found to be 0.06 ± 0.003 and 0.06 ± 0.003 μmol min^−1^ mg^−1^ protein, for Mace and EGA2248, respectively (Figure 3d). Heat treatment did not have a significant impact on Complex II activity, with the rate remaining at 0.07 ± 0.003 μmol min^−1^ mg^−1^ protein^−1^ in Mace, and 0.06 ± 0.002 μmol min^−1^ mg^−1^ protein in EGA2248. These rates refer to the electron transport through Complex II to UQ (Jones et al. 2013), and are 2500-10 000 fold lower than Complex II specific activity measured as the rate of succinate oxidation at the FAD redox centre (Huang et al. 2013).

The activity of Complex III in the two wheat varieties was not significantly impacted by treatment (Figure 3e). Rather, the specific activity differs by wheat cultivar. In Mace, the enzyme activity was determined to be 0.02 ± 0.009 and 0.03 ± 0.002 μmol min^−1^ mg^−1^ protein, at OT and HT, respectively. In EGA2248, the activity was shown to be 0.06 ± 0.006 and 0.04 ± 0.008 μmol min^−1^ mg^−1^ protein. Although the enzyme activity was not significantly altered by treatment temperature, the activity was significantly different between the two wheat varieties, and higher in the heat-sensitive variety. Similarly, the specific activity of Complex IV was determined to be largely unaffected by temperature (Figure 3f). The activity in Mace was determined to be 0.24 ± 0.04 and 0.16 ± 0.03 μmol min^−1^ mg^−1^ protein^−1^, at OT and HT. As with the findings for Complex III, the specific activity of Complex IV in EGA2248 was significantly greater than in Mace, at 0.37 ± 0.03 and 0.35 ± 0.06 μmol min^−1^ mg^−1^ protein. The number of copies of Complex III monomers, Complex IV, and cytochrome *c* (cyt *c*) in Arabidopsis is such that the stoichiometry supports the Complex III dimer-Complex IV supercomplex with a 1:1 ratio with cyt *c* (Fuchs et al. 2020). In fact, the flow of electrons between the two complexes is dependent on the equilibration time of cyt *c* in the intermembrane space, so there is a kinetic advantage to supercomplex assembly of Complex III and IV (Maldonado et al. 2021). The higher specific activity of both Complex III and Complex IV in EGA2248 is necessary to balance cyt *c* cycling. The common factor for these is cyt *c*, so it is reasonable to propose that, if it is not complex abundance, a possible explanation may be a difference in the amino acid sequence, which either weakens the interaction with cyt *c*, allowing for more rapid docking and undocking, or in the positioning of the redox centres that allow for a more rapid electron transfer (Solmaz et al. 2008, Xia et al. 2013, Maldonado et al. 2021). Unfortunately, the genome sequence of EGA2248 is not currently available to confirm this. Additionally, the cyt *c* substrate used in the assay from a mammalian source. The sequence similarity between cyt c from *Bos taurus* used in the current study, and that of *Triticum aestivum* was determined by Clustal Omega (https://www.ebi.ac.uk/Tools/msa/clustalo/) as 64%. It then stands to reason that the interaction of the mammalian cyt *c* with the binding site on Complex III will differ to its plant counterpart. It is then also possible that the activity difference observed here is limited to an *in vitro* effect due the affinity of the binding site for a bovine cyt *c* and the reaction conditions used.

To determine whether the differences in the specific activities of the individual complexes were caused by a change in complex abundance, representative peptides of subunits of Complexes I, II, III, IV, ATP synthase and type I and II NADH dehydrogenases were detected by selected reaction monitoring (SRM) mass spectrometry. The selected targets from a pre-existing in-house dataset were converted to current accession IDs, resulting in a dataset containing 275 transitions representing 84 peptides (Supplementary Table 4). Log2 fold change analysis between OT and HT is 0 for the majority of features studied in both wheat varieties, indicating that expression of peptides remains unchanged (Figure 3g and h). Although some features appeared to change in abundance, Student t-test indicated that these were not significant. One-way ANOVA with Tukeys Honestly Significant Difference (HSD) test indicated no significant differences in the relative abundance of peptides at 5 % significance level between the two wheat varieties. These results indicate that the loss of Complex I activity in the heat-sensitive variety is not from a decrease in complex abundance, and it then stands to reason that the effect lies in the membrane lipid remodelling.

Previous research on Complex I revealed a lipid requirement during purification to retain oxidoreductase activity (Sharpley et al. 2006). The changes to the lipid profile of the mitochondria of EGA2248 following HT could thus impact on the assembly, and the activity of the complex. BN-PAGE separates proteins under non-denaturing conditions, separating the respiratory complexes in their native state and with their associated subunits, and, with in-gel histochemical staining, detects the active complex (Sabar et al. 2005, Gazizova et al. 2020). This method was employed to assess if the assembly of the thermolabile Complex I is impacted during HT, which could explain the decrease in the specific activity. Three bands of high molecular mass were detected (Figure 3i), which align with activity bands detected by Gazazivo et al. (2020) for the supercomplexes incorporating Complex I from wheat root mitochondria. The relative intensity of the upper most band in EGA2248, corresponding to the supercomplex of I-III_2_-IV, decreased to 79% following HT. In comparison, the intensity in Mace did not decrease. The band representing supercomplex I-III_2_ has the highest intensity, as is expected given that the supercomplex I-III_2_ is known to be highly stable and the most abundant of the plant respiratory supercomplexes (Gazizova et al. 2020). The band intensity of Complex I in EGA2248 following HT was visibly less intense, and estimated to be ~86% of OT. This decrease in magnitude is observed in the specific activity measurement of Complex I in EGA2248 following HT. The Coomassie stained gel presented alongside its histochemically stained counterpart shows a slight decrease in the intensity of Complex I in EGA2248 following HT. This potentially points to an impairment in the assembly of the Complex I itself in EGA2248 following HT, or a dissociation of the subunits.

### Loss of Complex I activity in EGA2248 exacerbates mitochondrial redox stress

The mitochondrial tricarboxylic acid cycle (TCA) is a source of reducing equivalents that influences the redox state of the organelle. Oxidation of TCA cycle intermediates isocitrate and malate yields reducing compound NADH, which is a substrate for the ETC at Complex I (Sweetlove et al. 2010). We undertook quantitation of TCA cycle metabolites to assess the impact of Complex I activity impairment on the TCA cycle, and thus the redox state of the organelle.

The abundance of pyruvate and α-ketoglutarate in Mace did not change significantly during HT from 1.27 ± 0.12 to 1.90 ± 0.26, and 7.43 ± 0.78 to 8.80 ± 0.92, respectively (Figure 4a and d). The abundance of these metabolites was not significantly different in EGA2248, and similarly they did not increase significantly during HT, with pyruvate ranging from 0.99 ± 0.16 at OT to 0.70 ± 0.11 pmol mg^−1^ fresh weight at HT, and α-ketoglutarate measured at 9.16 ± 1.26 at OT to 6.66 ± 0.73 pmol mg^−1^ fresh weight at HT.

**Figure 4.**
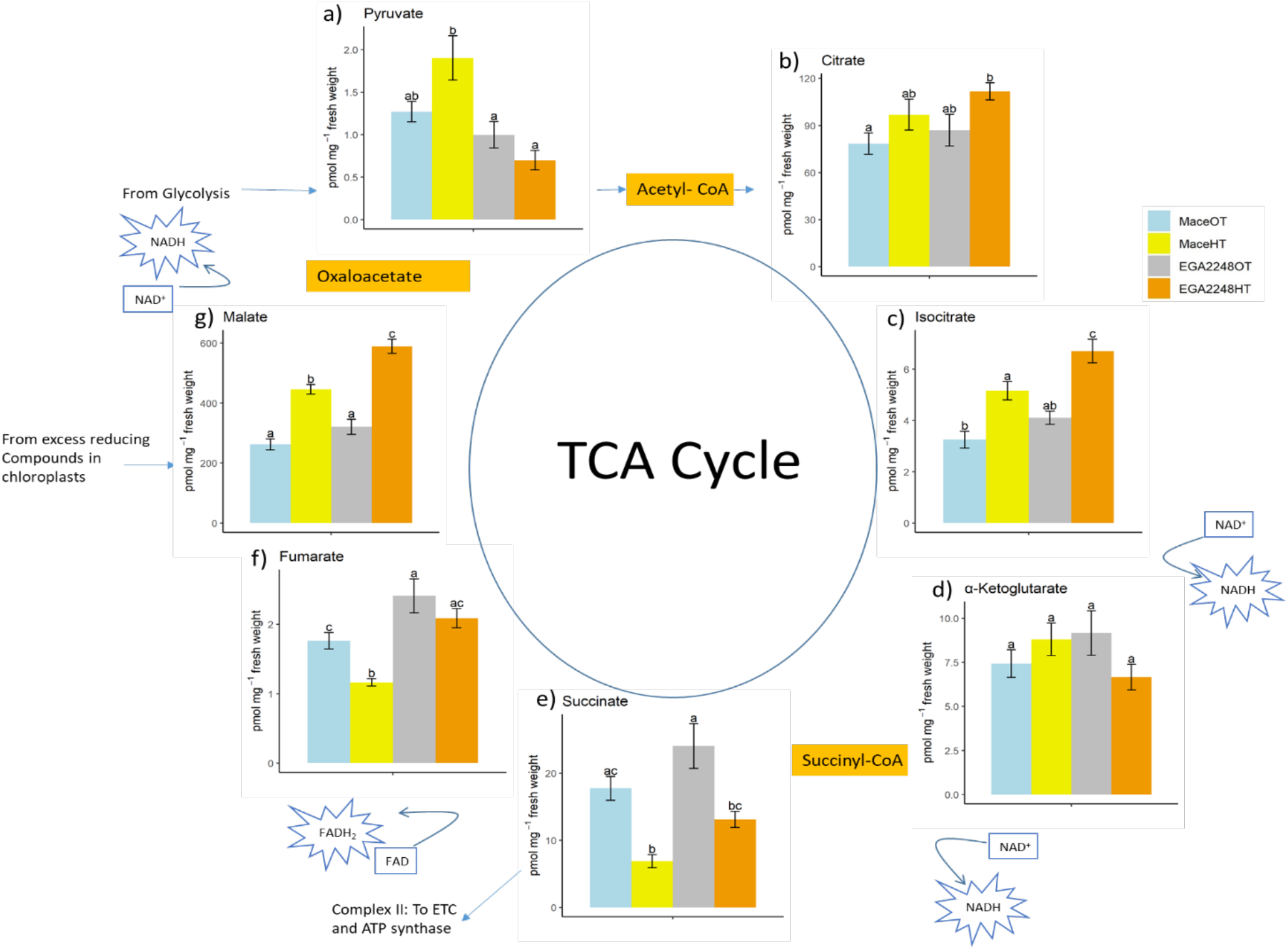
Absolute quantitation of detected TCA cycle metabolites **a)** Pyruvate, **b)** Citrate, **c)** Isocitrate, **d)**α-Ketoglutarate, **e)** Succinate, **f)** Fumarate, and **g)** Malate, from whole tissue obtained from OT and HT treated wheat seedlings, as pmol of metabolite mg^−1^ fresh weight. (**a-g;** two-way ANOVA with repeated measures p ≤ 0.05). Values are the mean of n≥ 8 biological replicates (± SEM).

Isocitrate (Figure 4c) was found to increase significantly, from 3.25 ± 0.33 to 5.15 ± 0.36 56 pmol mg^−1^ fresh weight in Mace, and 4.10 ± 0.25 to 6.71 ± 0.46 56 pmol mg^−1^ fresh weight in EGA2248, while in contrast, succinate (Figure 4e) and fumarate (Figure 4f) decreased in abundance. Isocitrate dehydrogenase converts isocitrate to α-ketoglutarate and the reducing equivalent NAD(P)H (Popova et al. 1998). The activity of the enzyme is competitively inhibited by increasing NADH, and non-competitively inhibited by NADPH, a reducing equivalent generated during photosynthesis (McIntosh et al. 1992). This feedback mechanism makes isocitrate dehydrogenase a regulatory step of the TCA cycle, to limit the production of further reducing equivalents (Popova et al. 1998). The results show an accumulation of isocitrate under HT, and this acts as the limiting step in the TCA cycle. The concurrent decrease in succinate and fumarate suggests that the TCA cycle is modified during heat stress and that energy production decreases when the ETC is compromised. As the abundance of α-ketoglutarate did not increase, the inhibition of isocitrate dehydrogenase is likely a result excess reducing power.

The most abundant metabolite measured by this approach, malate, increased significantly in Mace from 262.51 ± 18.69 to 445.89 ± 16.16 pmol mg^−1^ fresh weight, and from 320.82 ± 25.12 to 589.35 ± 23.56 pmol mg^−1^ fresh weight in EGA2248 (Figure 4g). The abundance of malate is at least 100-fold higher than its precursor of the TCA cycle, fumarate. The significant increase of key metabolites isocitrate and malate in the heat-sensitive wheat cultivar indicates that the loss of Complex I activity makes the mitochondria less-effective in dissipating the excess reducing equivalents, and causes redox stress.

## Discussion

Mitochondrial membranes are a complex mixture of lipids, and the close association of respiratory protein complexes with the phospholipid bilayer means any perturbations in the membrane ultimately impact on the ATP synthesis required for wheat growth and development. For Australian wheat cultivars Mace and EGA2248, their differences in heat tolerance with respect to grain formation is also apparent by their rate of biomass accumulation at the vegetative stage, which is significantly lower in the heat-sensitive cultivar EGA2248 compared to the heat-tolerant cultivar, Mace. These findings support the previous reports by Collins (2015) that suggest that a vegetative growth response following heat shock can be used to predict the grain filling response. It then stands to reason that heat waves at the seedling stage limit the carbon assimilates that can be drawn upon later during development, namely during the energy expensive stage of grain filling.

Quantitation of the mitochondrial membrane lipids from wheat seedlings identified 70 lipid species whose abundance changed significantly and revealed a trend towards shorter acyl-tail chain length following heat shock. Only in the heat tolerant wheat variety Mace, was there a significant decrease in PE species 40:5 and 40:6. A similar result was previously observed when cv. Karl-92 and cv. Ventnor wheat were exposed to heat, with a significant decrease of PE in the heat-tolerant variety Ventnor (Narayanan et al. 2016). This mechanism may stabilize the bilayer structure of membranes, as the propensity for H_II_ phase transition increases with the lengthening of the acyl tails, whereas shorter acyl tails favour the bilayer form (Koynova et al. 1994, de Kroon et al. 2013).

A significant increase in PI during heat treatment was measured in both wheat varieties and was attributed to an increase in shorter chain 32:0, 34:1, 34:2 and 34:3 species. As PI acts as a structural component of membranes with a propensity for the bilayer form (Osman et al. 2011), it is possible that this remodelling preserves the bilayer structure under temperature stress. Additionally, the phosphorylation and hydrolysis of PI provides a reservoir of secondary signalling molecules which are widely reported to partake in stress-response signalling (Löfke et al. 2008). Similar to the wheat reproductive phase, rice spikelet fertility is temperature-sensitive and Wada et al. (2020) detected an increase in PI 34:3 synthesis in rice pollen, which the researchers concluded serves as a precursor for secondary signalling molecules which are necessary in activating the pathways that support spikelet fertility under these conditions. The results here endorse the view that the synthesis of PI increases during heat stress and may act as a source of signalling molecules to activate heat stress pathways.

The propensity towards shorter fatty acyl tails has further impacts on protein incorporation into the membrane, with shorter acyl-tail chain length decreasing phospholipid-phospholipid interactions and enhancing protein incorporation into membranes (McMullen et al. 1997). Protein incorporation into membranes relies on hydrophobic matching between the acyl-tails and the transmembrane domain of the protein (Bogdanov et al. 1999), and the modification of acyl groups may alter the structure and positioning of membrane proteins. Furthermore, the length and saturation of the acyl side chain impacts on membrane thickness (Li et al. 2012). Membrane thickness has been reported to affect the function of certain membrane proteins, including ATP-dependant ion pumps (Na^+^/K^+^ ATPase, Ca^2+^ Mg^2+)^ (Johannsson et al. 1981). Thus, the shortening of acyl tails may impact on respiratory complex activity.

Odd-chained fatty acids, which arise following α-oxidation of fatty acyls, have been detected at low abundances in all organisms, including plants (Harwood 1997, Rezanka et al. 2009). The identification of a pathogen-inducible oxygenase from tobacco leaves implies that α-oxidation plays a role in plant pathogen defense (Hamberg et al. 1999). Alpine plants, which are subject to large daily fluctuations in temperature, have been shown to contain odd-chain fatty acids, which the authors suggest are an adaptation of the membranes to withstand the extreme conditions, and this notion is supported by the observed increase in PE 35:2, 35:3 and 35:3 that was unique to the heat-tolerant wheat variety. The increase in the expression of phospholipids containing odd-chain fatty acyl group may indicate the induction of a stress response rather than a marker of the stress condition itself, with oxidized lipids acting as lipid-mediators with the potential to induce the expression of genes and enzymes used in plant defense systems (Hamberg et al. 1999, Tsydendambaev et al. 2004). Odd-chain containing phospholipids PE 33:3, PC 33:3, PI 33:3 and 35:3 were observed in wheat grown at 12 days of high temperatures, with the researchers reasoning that the co-expression is explained by a shared metabolic pathways involving α-oxidation of fatty acyls, (Narayanan et al. 2016). The results obtained here are in support of the proposed theory of Narayanan et al. (2016) that a shared metabolic pathway explains the co-occurrence of odd-chain fatty acyl during HT.

A key difference in the lipid remodelling between Mace and EGA2248 is the significant increase in phosphatidylserine (PS) in the heat tolerant variety during heat stress. As such, this notable feature may shed light onto the molecular mechanism of heat tolerance with respect to the lipo-protein network of the mitochondria. PS is structurally related to PE, and one well conserved pathway of PE synthesis is PS decarboxylation by PS decarboxylase at the inner mitochondrial membranes. PS decarboxylation is the primary source of PE in yeast, and this pathway also exists in plants (Bürgermeister et al. 2004, Nerlich et al. 2007). The accumulation of longer chained species, taken together with the apparent trend for a reduction in longer acyl chain PE, can be interpreted as an interference in the synthesis of PE species with a propensity for bilayer-destabilization and thus act as a mechanism for bilayer preservation under temperature stress conditions.

Phospholipid headgroups are able to recruit and bind peripheral membrane proteins, and the selective interaction of heat shock proteins (HSP) with PS has been documented. For example in plants, small heat shock proteins (sHSP) have been reported to convey thermotolerance in isolated mitochondria (Chou et al. 1989). The sHSP, HSP17 in cyanobacteria rigidifies membranes at high temperature, and also increases the transition temperature for phase transition of PE, and the interaction is mediated through the PS and PE headgroups (Tsvetkova et al. 2002). Small heat shock proteins have also been reported to protect Complex I during abiotic stress (Downs et al. 1998, Hamilton et al. 2001). For example, during salinity stress in maize, the exogenous addition of sHSPs to mitochondria was able to restore electron transfer through Complex I by 33% when compared to the rates without sHSP treatment (Hamilton et al. 2001). Also, the inactivation of sHSPs in murine cells subject to heat stress decreased both the electron transfer to ubiquinone, and NADH oxidation activity of Complex I (Downs et al. 1998). In the research presented here, the total quantity of PS increased significantly in Mace during heat stress. This may indicate that the increase in the total amount of PS in the heat-tolerant variety facilitates the recruitment and binding of sHSP to membranes to stabilise them during heat stress, and in addition to protect the activity of Complex I.

The presented results point to a mechanism whereby the heat-tolerant wheat variety, Mace, more effectively undergoes mitochondrial membrane lipid remodelling, to preserve the activity of Complex I. This facilitates the NADH cycling within the TCA cycle, avoiding the build-up of excess reducing power in the mitochondria, ROS generation and their accompanying pathologies. In contrast, the heat-sensitive variety EGA2248 experienced a decrease in Complex I activity during heat shock, and biomass accumulation was negatively impacted to a greater extent than that of Mace. To date, the role of PS in abiotic stresses are conflicting. For example the accumulation of PS is associated with early senescence in rice (Rani et al. 2020), while PS is increased in salt-tolerant sweet potato leaf lipidome under salinity, and exogenously applied PS is able to delay salinity-induced senescence in the leaves of a salt-sensitive variety of sweet potato (Yu et al. 2019). Similarly, the gene encoding a PS synthase involved in PS synthesis through the condensation of CDP-DAG with serine increases in expression in wheat roots in response to increasing aluminium toxicity and confers aluminium resistance when the gene is expressed in some yeast strains, yet results in necrotic leaf lesions when expressed in Arabidopsis (Delhaize et al. 1999). Thus, the precise function of PS in abiotic stress is yet to be fully elucidated, however it appears that it plays a crucial role in maintaining Complex 1 activity in wheat mitochondria during heat exposure, and is a potential future avenue for increasing the heat tolerance of wheat.

## Data availability

All mass spectrometry data is available on request.

## Acknowledgements

We acknowledge and celebrate the First Australians on whose traditional land this research was undertaken, and pay our respect to the elders past, present and emerging. A.M. was funded by the Australian Government Research Training Program (RTP) Scholarship. The research was also generously supported by the Charles and Annie Neumann Fellowship PhD Scholarship in Agriculture, the Underwood PhD Completion Scholarship. The authors also thank for funding towards this project from the Research Grant Support Scheme from the Faculty of Science, University of Melbourne.

## Authors contributions

A.M. and N.L.T. conceived the research plans; N.L.T. and U.R. supervised the experiments; A.M. and T.R. performed the experiments; A.M. and N.L.T. wrote the article.

## Supplementary Information

**Supplementary Table 1.**

Specifications of phospholipid SRM transitions for phospholipid quantification from isolated mitochondria. The target list includes the phospholipid standards from the “Lipid Mix”, designated by the suffix “IS”.

**Supplementary Table 2.**

Specifications of phospholipid SRM transitions for cardiolipin quantitation from isolated mitochondria.

**Supplementary Table 3.**

Specification of peptide SRM transitions for mitochondrial electron transport chain proteins from isolated mitochondria. The list contained 275 transitions from 84 peptides.

**Supplementary Table 4.**

Conversion of Accession ID from the in-house dataset (TGAT 1.0) to current accession IDs (RefSeq 1.0) was by searching peptides against were updated by searching the IWGSC RefSeq 1.0 using https://wheatproteome.org/. For convenience, the lowest genome ortholog is deemed a protein match.

**Supplementary Table 5.**

The absolute quantification of the 135 phospholipids, as detected by SRM, and expressed as nmol lipid per mg of mitochondrial membrane protein (nmol mg-1 protein). The values are presented as the mean of 4 biological replicates, ±SEM.

